# BCN057, a Modulator of GSK3β, Induces KRAS G12D Mutant Pancreatic Cancer Cell Death

**DOI:** 10.1101/2021.09.03.458938

**Authors:** Elizabeth M. Singer, Rishi Man Chugh, Payel Bhanja, Adrian Gomez, Lucy Gao, Julian P. Whitelegge, William H. McBride, Subhrajit Saha, Andrew J. Norris

## Abstract

Effective treatment for Pancreatic Cancer remains a major challenge due to its resistance to radiation/chemotherapy and poor drug permeability. Moreover, treatment induced normal tissue toxicity, mainly to the duodenum and gastrointestinal epithelium, is common and is a dose limiting event, while toxicity to the pancreas is relatively rare^1–3^. Gastrointestinal toxicity, however, often results in interruption, reduction or premature withdrawal of anti-cancer therapy which is a very significant factor impacting the overall survival of patients being treated. Therefore, development of a therapeutic strategy to selectively sensitize tumor tissue without inducing normal tissue toxicity is important. In this manuscript, we show that the novel small molecule BCN057 can modulate chemo-sensitivity of oncogenic RAS pancreatic cancer cells while conversely protecting normal intestinal epithelium from off target toxicity. In particular, BCN 057 protects Lgr5 positive intestinal stem cells, thereby preserving barrier function. Further, it is demonstrated that BCN057 inhibits GSK3β and thereby induces a pro-apoptotic phosphorylation pattern on c-Jun in KRAS G12D mutant pancreatic cancer cells (Panc-1) leading to the restoration of PTEN expression and consequent apoptosis. This appears to be a new mechanistic observation for the oncogenic RAS phenotype. Lastly, concurrent with its GSK3β inhibition, BCN057 is a small molecule inhibitor of PD-1 expression on human T-lymphocytes co-cultured with human pancreatic cancer cells. In summary, BCN057 can promote synthetic lethality specifically to malignant cells and therefore should be considered to improve the therapeutic ratio in pancreatic and epithelial cancer treatment in conjunction with chemotherapy and radiation.

## Introduction

Pancreatic cancer remains one of the most lethal malignant neoplasms with projected 2021 estimates from the American Cancer Society of 60,430 people diagnosed and 48,220 people who will die. In the US, this amounts to about 7% of all cancer deaths, with a 5-year survival rate standing at only 9%, even though it is only 3% of the total cases. Globally, pancreatic cancer is the seventh leading cause of cancer-related deaths out of the over 450,000 reported annually^4^. Pancreatic cancer presents several unusual barriers to therapeutic intervention. The small intestine is in front of the pancreas and in the path of radiation used to treat pancreatic cancer, along with the duodenum. Because of this, severe epithelial damage limits treatment dose ^5^. Chemotherapy further exasperates the epithelial tissue. Today, there are no drugs to prevent the epithelial toxicity from radiation and chemotherapy which is a principal factor determining maximum tolerated dose (MTD), with a consequent negative impact on tumor control. The small molecule BCN057 was originally designed under the NIH initiative for drugs to counter normal tissue damage from radiation (nuclear threat). BCN057 is a novel small molecule which inhibits GSK3β activity promoting survival and self-renewal of the LGR5+ intestinal stem cells in the tissue, but it has also been shown to cause apoptosis in pancreatic and other cancer cells possessing oncogenic KRAS mutations^6^. Radiation and chemotherapy draw their specificity by targeting cells which are rapidly dividing, including cancer cells and active stem cells. GI epithelial tissue regenerates every 4-5 days in healthy adults from a highly proliferative stem/progenitor cell compartment^7,8 9^ and thus, this organ presents particular off target challenges for many therapies. The tissue response to radiation and chemotherapy involves apoptosis within this compartment^10^ with the consequence that tissue tolerance and regeneration are severely compromised. Our previous report showed that BCN057 induces β-Catenin localization with consequent TCF/LEF gene expression within normal LGR5+ stem cells, thus, preventing normal epithelial tissue damage from radiation^6^ and here, chemotherapy (5FU). We have discovered that GSK3β inhibition at S-9 also restores apoptosis in KRAS mutant pancreatic cancer cells through a c-Jun-dependent mechanism. Over 70 cancer models have been looked at with BCN057 both at The National Cancer Institute (NCI), The Armed Forces Radiobiology Research Institute (AFRRI) and academic institutions that along with this data indicate that BCN057 has antineoplastic activity and works in synergy with radiation and chemotherapy. BCN057 is a breakthrough for epithelial cancers and pancreatic cancer treatment and may eventually have significant implications for other areas (oral mucositis, proctitis, radiation enteritis or even pediatric radiation oncology). Pancreatic cancer is the third-leading cause of cancer-related death in the United States and has been projected to continue its increase in number in the next several decades, unlike colon, lung, breast and prostate cancers^11^. Radiation and chemotherapy are the only options for non resectable or borderline resectable pancreatic cancer; however, treatment options severely compromise normal tissue toxicity primarily in the duodenum. The treatment resistant nature of these tumors and the epithelial toxicity significantly reduces the treatment efficacy. Sensitizing the tumor tissue and/or reducing the normal tissue toxicity is needed because both impact overall survival and progression free survival.

Here we show that BCN057 causes apoptosis in KRAS mutant pancreatic tumor cells and simultaneously promotes epithelial cell regeneration in small bowel damaged by chemotherapy and radiation by preserving normal Lgr5+ stem cells. In addition, we show it is a potent inhibitor of PD-1, making it a possible immune checkpoint (IO) inhibitor.

## Results

### BCN057 inhibits pancreatic cancer cell growth by causing apoptosis

Previously we reported the anti-neoplastic effect of BCN057 in over 60 cancer cell lines^6^ which include inhibiting Renal Cancer: A498, Leukemia: RPMI-8226, CCRF-CEM, Ovarian Cancer: OVCAR-5, Breast Cancer: MDA-MB-231, which possess mutations in KRAS^12^. To determine the effect of BCN057 in pancreatic cancer, Panc-1 cells, a G12D KRAS mutant cell line^13^ were treated with BCN057 for 48 hours. ATPlite™ assay (PerkinElmer) an assay which measures cell viability, was used to demonstrate a significant decrease in PANC-1 cells viability with increasing doses of BCN057 (Figure 1).

**Figure 1:**
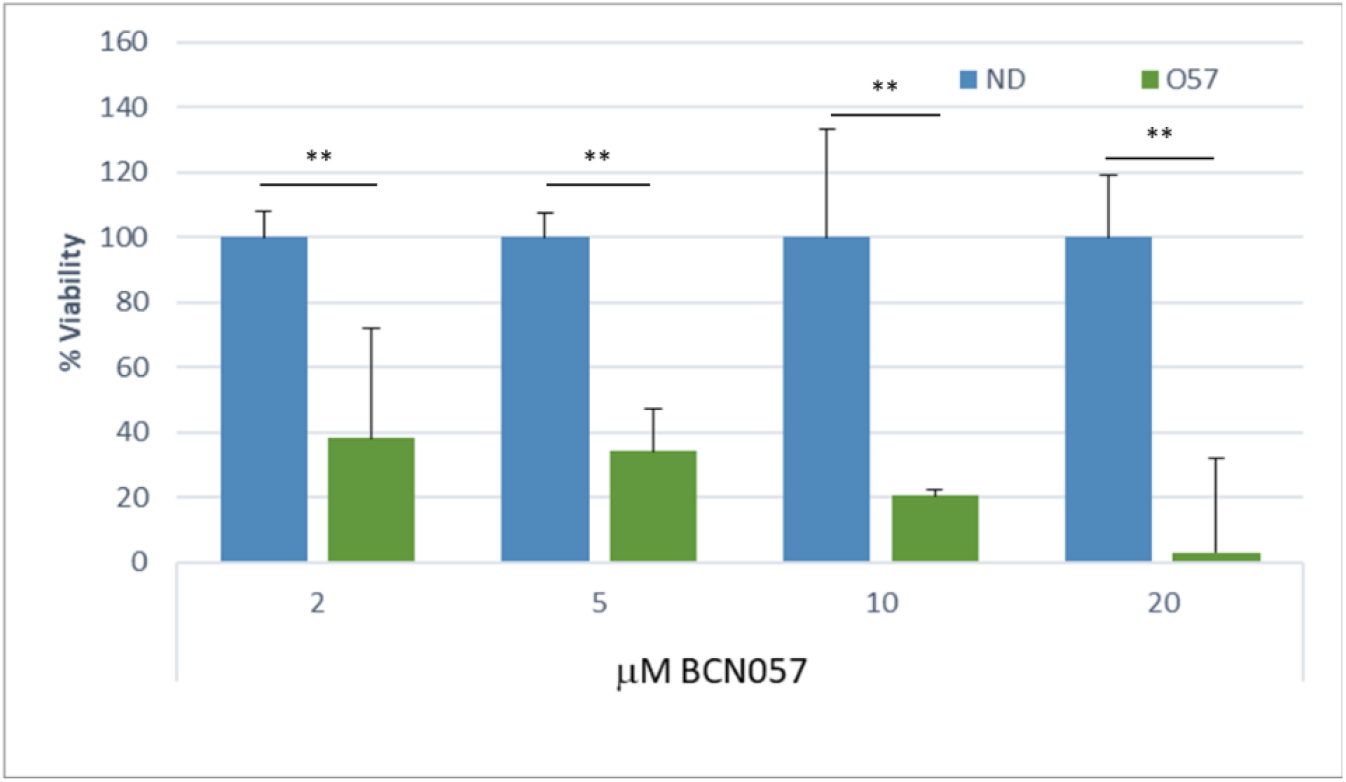
Effects of BCN057 on the survival of pancreatic cancer cell lines as assessed by ATP Lite™ assay (*Perkin Elmer*) with incubation 72 hrs post application of drug. All data points are n=5. Control (ND) received no drug and BCN057 at 2, 5, 10, and 20 µM final concentration. The Percent viability was calculated by dividing the average value for each condition by the average value of the control cells (V) at each time point. *p<0.05 **p<0.005.

This data (Figure 1) was assessed with Quest Graph™ IC_50_ Calculator^14^ using Equation 1 which yielded an IC_50_ of 1.33 μm. This value compares with other analysis of cytotoxic drugs, such as gemcitabine, on pancreatic ductal adenocarcinoma cells (AsPC-1, BxPC-3, MIA PaCa-2 and Panc-1), where IC_50’s_ ranges from 494 nM to 23.9 μM^15^.

The drug was found to show cytotoxicity toward the cells at all concentrations in the dose response curve. A dose proportionality study was conducted in C57Bl/6 mice and these results indicate that the IC_50_ values are well within the range of concentrations found in plasma during efficacy studies in rodents (Supplementary data Figure 1).

Because Panc-1 cells possess a heterozygous missense mutation in codon 12 (p.G12D; GGT > GAT of K-RAS) similar to other KRAS mutant tumor cells^13^, the phenotype has lost the capacity for apoptosis and is consequently chemo-resistant. BCN057 treatment appears to rapidly and dose dependently induce apoptosis as early as 20 min of exposure (Figure 2).

**Figure 2:**
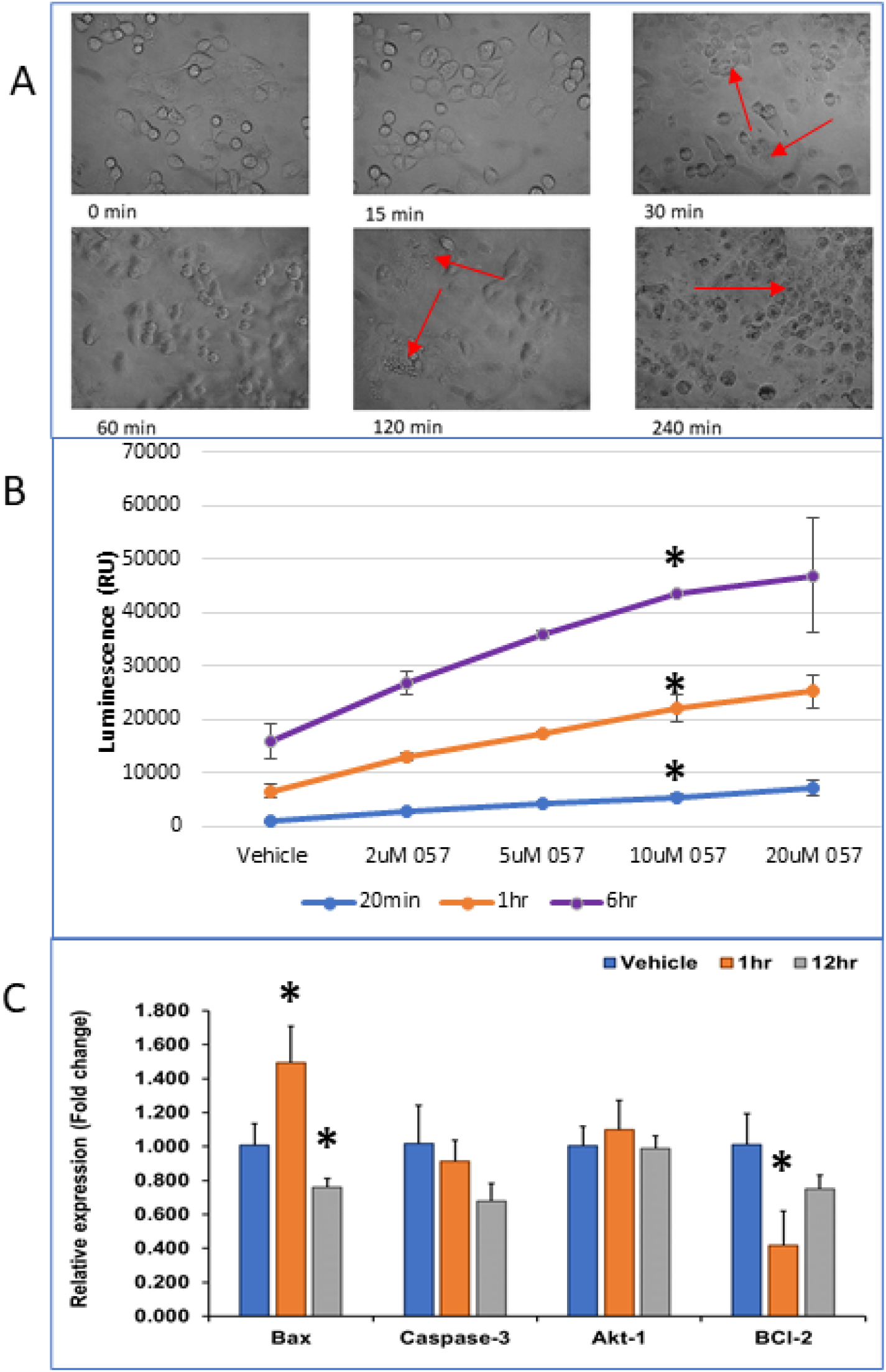
**A**. Time lapse of pancreatic cancer cell growth after treatment with 10 µM BCN057. Blebbing and cell fragmentation was noted within 20 mins of incubation with drug. Cell fragmentation is seen between 20-30 mins after exposure to the drug. **B**. Annexin V Apoptosis and Necrosis Assay in Panc-1 cells: Panc-1 cells were exposed to serial dilutions of BCN057 using the RealTime-Glo™ Annexin V Apoptosis and Necrosis Assay Reagent. The plate was incubated at 37 °C, 5% CO_2_ and luminescence (Annexin V binding) was measured over a period of 6 hrs with each data point run in duplicate. No change was observed in fluorescence over the same time period. Apoptosis appeared to increase with increasing concentrations of BCN057. Apoptosis was observed in a dose dependent fashion and time dependent with time points at 10 mins, 30 mins, 1 hrs. and 6 hrs. p values were assessed for 10 μM only as this concentration was frequently used in other assays. **C**. qPCR analysis of pro apoptosis and anti-apoptosis markers in Panc-1 cells. BAX is significantly elevated, and BCl-2 is decreased 1 hr. after exposure to BCN057 *p<0.05. Red arrows indicate apoptotic bodies emerging starting 20-30 mins.

Subsequent microscope images showed cell blebbing and fragmentation indicating possible apoptosis (Figure 2A). To validate the involvement of apoptosis in BCN057 induced cell death, the PANC-1 cells with/without BCN057 were subjected to Annexin V staining. A dose and time dependent increase in apoptosis was detected up to 6 hours later in PANC-1 cells treated with BCN057 (Figure 2B).

Further qPCR analysis was performed on Bax, BCl-2, Akt-1 and Caspase-3 which demonstrated that BCN057 induces expression of the pro-apoptotic gene Bax but inhibited anti-apoptotic Bcl2 gene expression in PANC-1 cells at 1-hour post treatment (Figure 2C). The expression profile did not affect Akt-1 or Caspase-3, consistent with caspase-independent cell death^16^.

### BCN057 inhibits GSK3β signaling to restore apoptosis in PANC-1 cells

As described, the PANC-1 cell line possesses a KRAS G12D mutation and has lost the capacity for apoptosis^17^,^18^ which is a hallmark of oncogenic RAS mutations. To investigate the role BCN057 may have in the restoration of apoptosis in this cell line, the GSK3β- c-Jun relationship was assessed, as this has been directly linked to apoptosis^19^. Others have reported a link with GSK3β and c-Jun^20^ and apoptosis along with the fact that GSK3β is over expressed in the oncogenic KRAS phenotype^21^. Analysis of PANC-1 cells for GSK3β inhibition over time after exposure to 10 µM BCN057 showed rapid induction of phosphorylation at S-9 (Figure 3 A-B). This inhibition was also noted in other non-transformed cell types such as HEK293 (Supplementary data, Figure 2). Phosphorylation at T93 in c-Jun was measured as a function of total c-Jun. Significant phosphorylation at T91/93 was observed in an immunoblot experiment which has been reported to be associated with the activation of c-Jun’s pro-apoptotic activity^22,23^. This activity was not induced in non-transformed cells when treated with BCN057^6^, even non-transformed HEK298 epithelial cells (Supplementary data, Figure 3).

**Figure 3:**
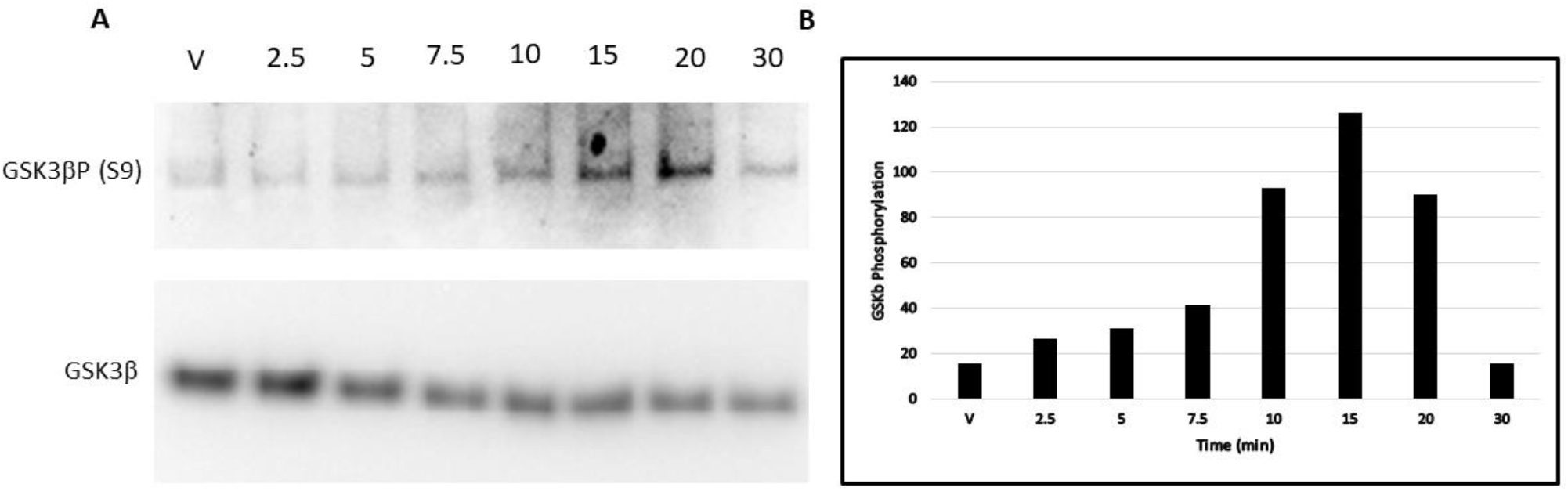
**A**. GSK3β phosphorylation at S-9 Increases following the addition of 10 µM BCN057 in Panc-1 Cells. **B**. Each value is relative expressed as a ratio of phosphorylated GSK3 β /total GSK3 β protein.

In KRAS dependent tumors, c-Jun acts to suppress phosphatase and tensin homolog (PTEN) expression at the promoter level, which is essential for anti-apoptosis and cellular transformation by oncogenic RAS^24^. Thus, we investigated PTEN protein expression levels in response to BCN057, and found it increased up to 1 hr (Figure 4B-C), with trailing decrease in the apoptotic phase thereafter, consistent with our hypothesis (Figure 2).

**Figure 4:**
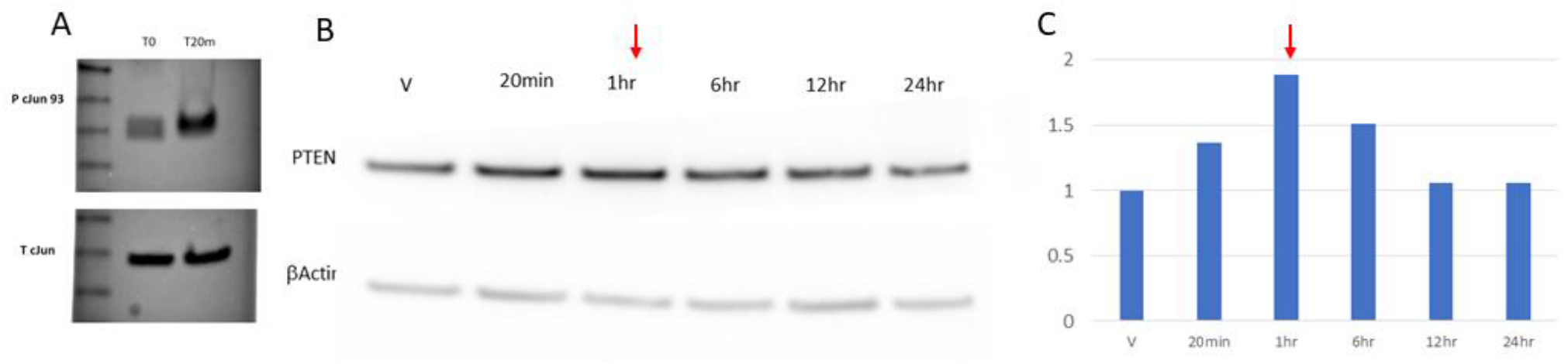
**A**. Phospho-c-Jun (T93): BCN057 significantly increases JNK dependent phosphorylation in KRAS mutant Panc-1 cancer cells when 20 mins after drug treatment. Phosphorylation of c-Jun at T91/93 is known to induce apoptosis. Total c-Jun protein level was unchanged. **B**: PTEN expression in Panc-1 cells. Western blot of PTEN protein expression over 24 hrs. with βActin stain from the same sample with vehicle or BCN057 treatment (10 µM) at the indicated time points. **C**. Histogram of the ratio of PTEN to βActin densitometry showing the fold increase in PTEN level in **B**. Red arrows indicate the onset of apoptosis.

### BCN057 augments anti-neoplastic effect of chemotherapeutic agents

The anti-apoptotic phenotype of KRAS is associated with significant chemo-resistance. To examine whether BCN057 enhances the cytotoxicity of the chemotherapeutic agents 5-Fluorouracil(5FU), PANC-1 cells were pre-treated with 5FU as reported^25^ and incubated with BCN057. A combination of 5FU and BCN057 at equal concentrations and conditions showed greater cytotoxicity than 5FU or BCN057 alone (Figure 5A). The effect followed a dose dependent response for BCN057 and was consistent over 48 hrs. after BN057 treatment (24 and 72 hrs. yielded similar results, data not shown).

**Figure5:**
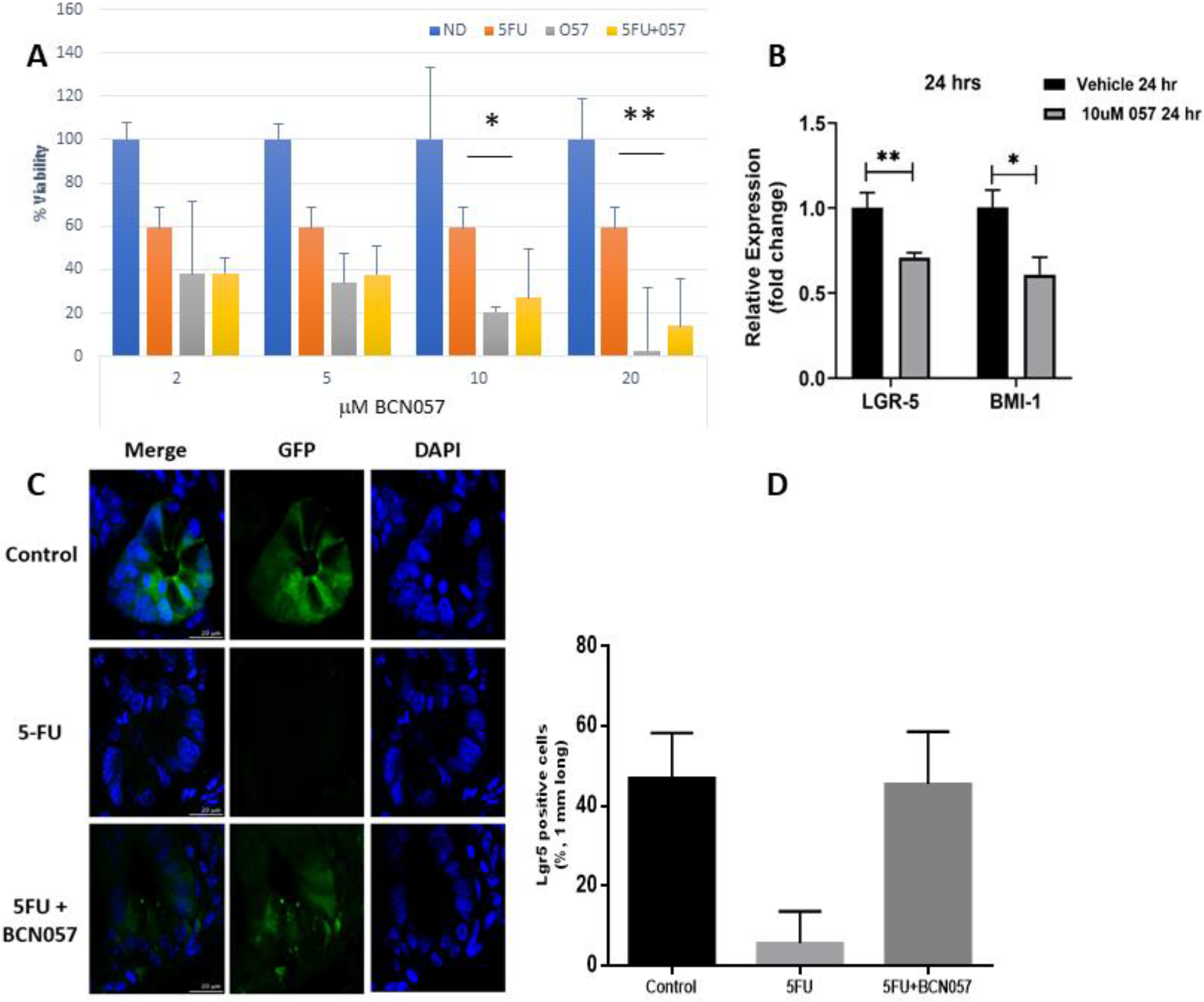
**A**. Effects of BCN057 and 5-FU on the survival of pancreatic cancer cell lines as assessed by ATP Lite™ assay (*Perkin Elmer*) with incubation of all drugs at 48 hrs. All experiments were in duplicate with 5FU and triplicate with 5FU with BCN057. Control (ND) received no drugs. Study groups include 5FU (50 µM), 057 (BCN057 at 2, 5, 10, and 20 µM or 5FU + BCN057 (BCN057 at 2, 5, 10, and 20 µM in combination with 5FU at 50 µM). The Percent viability was calculated by dividing the average value for each condition by the average value of the control cells (V) with standard deviation at each time point. **B**. Relative LGR-5 and BMI-1 expression in Panc-1 cells treated with 10 µM BCN057. **C**. BCN057 protects GI epithelium and the LGR5 stem cells (green) from 5FU cytotoxicity. Stains of GI crypts with green = LGR5+ stem cells. Mice were treated with 5FU and 5FU+BCN057 for 5 days. BCN057 was administered for 2 extra days after 5FU treatment. Small intestine samples were harvested and stained for LGR5 at 3 days after the last 5FU injection. **D**. Quantitation by % of LGR5+ cells remaining in 1 mM sections at the crypt base showing prevention of 5FU induced apoptosis in LGR5+ stem cells in GI. *p<0.05, **p<0.005.

#### LGR5 and BMI1 in pancreatic cancer

LGR5 has wide-spread expression in pancreatic tumor cells^26^. Others have reported that BMI1 may be a key contributor to chemoresistance and invasiveness^27^ in Panc-1 so we tested if BCN057 would increase these stem cell markers in Panc-1 cells but found no significant increase in abundance (Figure 5B).

### Epithelial LGR5 Stem cells

Previous reports from our group as well as others demonstrated that Lgr5+ve crypt base columnar cells are sensitive to chemoradiation therapy^5^. We have also reported that protection of Lgr5+ve intestinal stem cells (ISCs) from treatment -induced toxicity is crucial to minimize intestinal epithelial damage^6^. 5FU is a component of FOLFIRINOX and the active agent metabolized from Capecitabine (Xeloda®). To examine the effect of BCN057 against chemo-induced toxicity in Lgr5+ ISCs, Lgr5-Cre-ERT-GFP mice were pre-treated with 5FU as reported^5^ and then subjected to BCN057 treatment. Confocal microscopic imaging of jejunal sections clearly showed significant loss of Lgr5+ ISCs in 5FU-treated mice. However, 5FU-treated mice receiving BCN057 retained a significant presence of Lgr5+ve ISCs (Figure 5 C-D) suggesting BCN057 protects ISCs from chemotoxicity.

#### Combined chemotherapy and cytotoxicity

Other cytotoxic drugs commonly used in the treatment of pancreatic or colorectal cancer were then examined in Panc-1 cells in combination with equivalent doses of BCN057 over 24, 48 and 72 hrs. In each case, apoptosis was induced with enhanced statistically significant sensitivity to the cytotoxicity of the drugs by the combination with BCN057 at all time points except one (Irinotecan (IRI) and oxaliplatin (Ox) at 72 hrs.) (Figure 6). This trend remained the same with a separate study using much higher doses of Oxaliplatin (75 µM), Irinotecan (50 µM) and 5FU (50 µM) with BCN057 remaining at 10 µM with n = 6 (data not shown). This is consistent with the observation of others^28^ where GSK-3β has been shown to regulate the sensitivity of MIA-PaCa-2, Pancreatic, and MCF-7 breast cancer cells to Chemotherapeutic Drugs.

**Figure 6:**
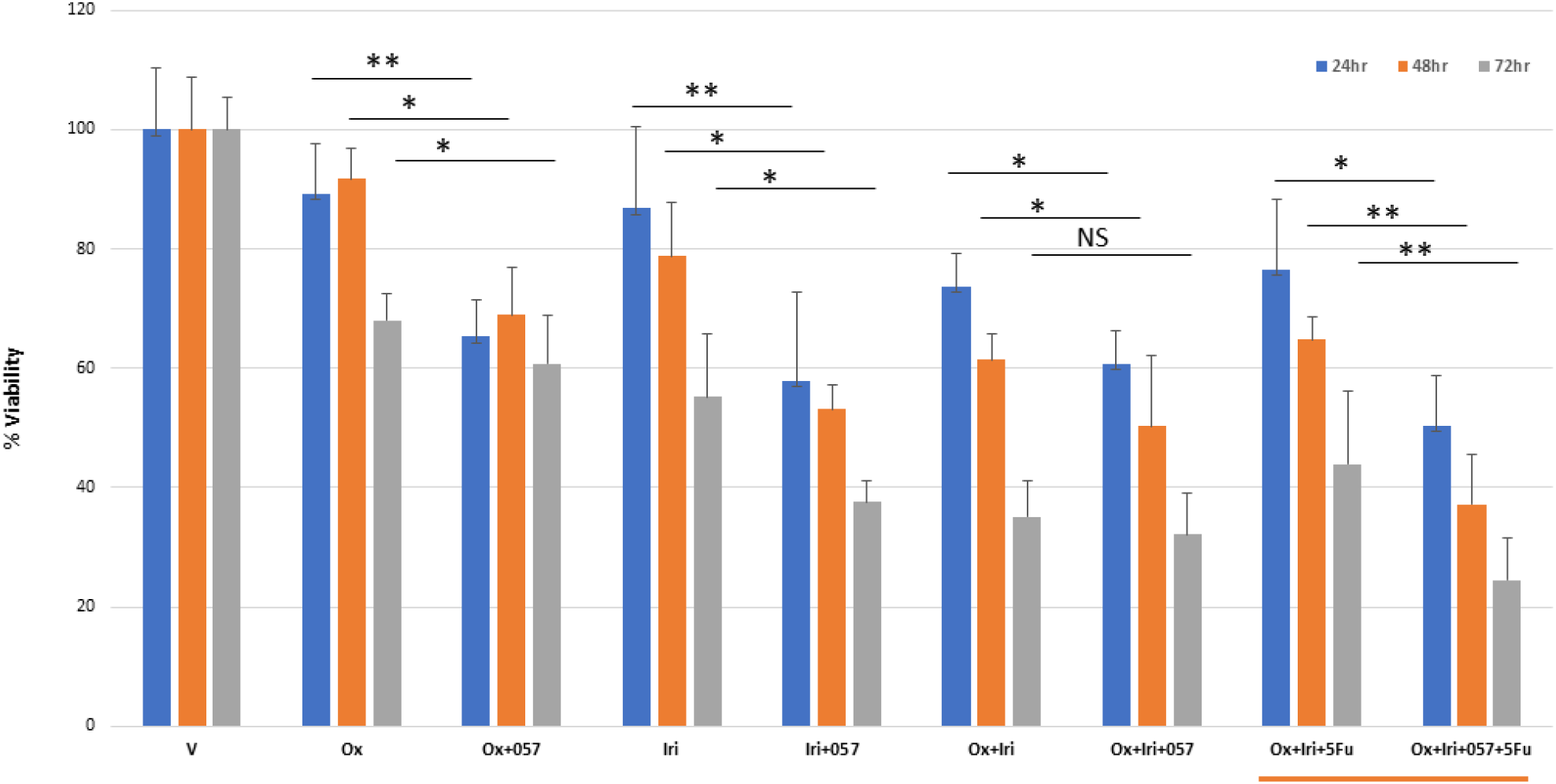
Cytotoxicity of Oxaliplatin and Irinotecan, individually or in combination with 5FU and BCN057 on Panc-1 cells. Cells were incubated with Oxaliplatin, Irinotecan and 5FU with concentrations at 10 µM each and BCN057 (057) also at 10 µM with n = 4 for all assays. In all cases, the addition of BCN057 resulted in a general trend of decreasing tumor cell viability over 72 hrs. *p<0.05 **p<0.005.

### BCN057 and Immune Checkpoint blockade

An emerging method of treatment for many forms of cancer, including PDAC, targets immune checkpoint inhibitors. Because GSK3β is a key regulator of immune checkpoints and in particular the PD-1/PDL-1 axis^29, 30, 31^ we sought to examine the effect of BCN057 on PD-1/PDL1 expression. A co-culture system consisting of PANC-1 cells and human T-cells separated by a membrane was used as previously described^32^. Panc-1 cells were co cultured with T-Cells for one day and subsequently treated with BCN057 at 10 µM for up to 12 hrs (Figure 7). PD-1 expression on human T-lymphocytes was decreased significantly within the 1^st^ hour and sustained over the 12 hrs post treatment as shown in Figure 7A. PDL-1 expression on Panc-1 cells was also decreased (Supplementary Figure 4), however, the cells were undergoing rapid apoptosis, as described, and the decrease is most likely due to this rather than a regulatory mechanism. In a separate study using mouse T-lymphocytes, 2 Gy radiation was used to induce apoptosis with increasing amounts of BCN057. The results demonstrated the preservation of the T-lymphocytes by BCN057 in the presence of radiation (Figure 7B) with an EC_50_ = 5.34 µM, thus establishing the difference in biological response between the tumor and immune cells.

**Figure 7:**
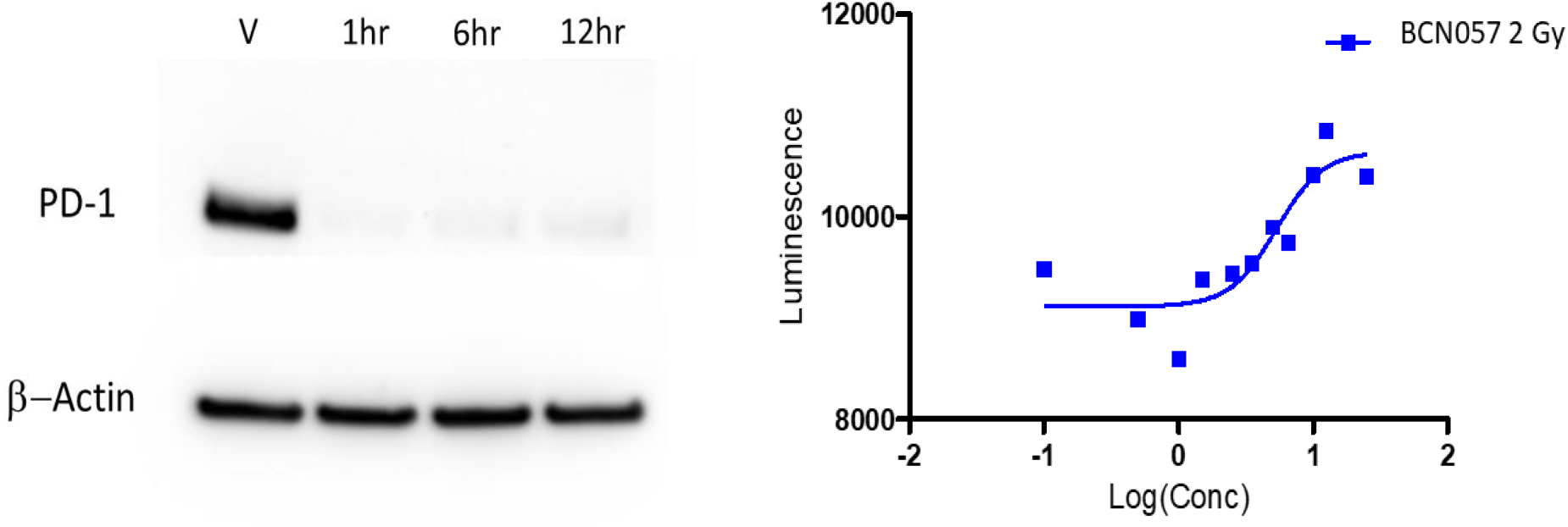
**A**. Western blot showing PD-1 expression on human T-lymphocytes co cultured with Panc-1 cells with no drug (vehicle) or 1, 6 and 12 hrs. with 10 μM BCN057. **B**. Inhibition of GSK3β induces survival of Lymphocytes when exposed to radiation. In vitro viability assay using increasing concentrations of BCN057 on irradiated mouse T-lymphocytes EC_50_=5.34μM which is within the plasma concentrations observed for the drug.

## Discussion

We reported previously that BCN057 significantly reduces tumor burden in a KRAS G13R mutant cancer and radiosensitizes colon cancer without promoting toxicity in small bowel^6^. Moreover, in vivo and in human ex vivo organoid models we have shown BCN057 radio protects intestinal stem cells and consequently, preserves barrier function. Here, we have shown that BCN057 modulates a tumoricidal effect by activating apoptotic cell death in the invasive KRAS G12D mutant cell line Panc-1. In a co-culture system consisting PANC-1 cells and TILs, BCN057 inhibits PD1 expression in T-lymphocytes. Moreover, in a mouse model of 5FU-mediated chemotoxicity^5^, BCN057 ameliorates chemo-induced intestinal damage, a major limitation in pancreatic cancer treatment.

In normal cells, BCN057 induces phosphorylation at S-9 in GSK3β located in the beta-catenin destruction complex. This leads to an accumulation of free non-phosphorylated beta-catenin in the cytosol, which translocates to the nucleus and trans-activates Wnt-target genes together with the T-cell factor (TCF)/lymphoid-enhancing factor (LEF) family of transcription factors. Thus, inhibition of GSK3β leads to a pharmacological activation of the downstream Wnt-signaling pathway in normal epithelial stem cells^6^. Given that this pathway is often ascribed to growth and survival, pharmacologic activation of the pathway in oncology, surprisingly, has not been well explored. GSK3 is reported to phosphorylate many pro-oncogenic targets in cells, including colorectal cancer, and concerns have been expressed that modulation of this pathway could induce cancer. However, there are no reports of cancer induction by agents that inhibit GSK3 signaling to date. Importantly, lithium (orotate, chloride or carbonate) that inhibits GSK3β and is a potent activator of canonical wnt signaling that has been widely used in humans since the 1950s throughout the world and the existing data show its use is not associated with an overall increased risk of cancer^33^. It is, however, associated with suppression of the metastatic potential of colon cancer cells^34^. Prior to this report, it has been demonstrated that prolonged inhibition of GSK3 β activates the c-Jun N-terminal kinase (JNK) pathway and triggers an apoptotic response in human pancreatic cancer cells but surprisingly not in human non-transformed pancreatic epithelial cells^35^. This is consistent with what we have observed with BCN057 in Panc-1 cells, and it is important to note that RAS activated cells are, in many respects, not wired the same as non transformed cells. Oncogenic RAS effectively inhibits PTEN expression via the RAF-MEK-ERK-c-Jun pathway, and thereby prevents apoptosis to promote cellular transformation^24^. This is a fundamental hallmark of the oncogenic RAS phenotype. In addition, mutant KRAS also increases GSK-3β gene expression^36^,^37^. Reports have also shown that in the oncogenic RAS phenotype, the AKT pathway is activated but without GSK3β phosphorylation at S-9^38^. This may be why GSK3β inhibition, a negative regulator of JNK, activates JNK^23^,^20^ in turn activating Jun, fos and c-Jun expression and phosphorylation. To our knowledge, we are the first to show that inhibition of GSK3β is correlated to a Thr (T93) Phosphorylation on c-Jun (likely by JNK1)^39^ with a consequent restoration of PTEN expression and apoptosis in oncogenic RAS cancer cells. Further, we are not aware of any other reports showing the mechanism of T93 phosphorylation by a drug to activate the c-jun pro-apoptotic feature in cancer, however this has been reported in neurobiology^19^,^23^. We have not observed the same T93 phosphorylation in non transformed cells in response to BCN057.

Further, the mechanism of cytotoxicity of BCN057 is independent of the nature of the standard of care chemotherapy such as Oxaliplatin, Irinotecan or 5-FU. As illustrated in Figure 1, 2, 5 and 6, there is a dose-dependent toxicity with BCN057 in Panc-1 cells, but this appears to improve the cytotoxicity of other chemotherapy or their combinations commonly used in pancreatic cancer treatment through the restoration of apoptosis. It was important to investigate this given the link with β-catenin and chemo-resistance. Others have also reported that GSK3 inhibition upregulates β-catenin and c-Myc resulting in the abrogation of KRAS-dependent tumors^40^.

A previous report has shown that LGR5 is widely expressed in all cells of pancreatic tumors^26^ and using a marker in beta cells hypothesized that cancer cells migrate from the islets to proliferate and contribute to the formation of cancerous ducts. During this process they cease insulin expression and activate LGR5 as a possible marker for pancreatic ductal adenocarcinoma (PDAC). Others have implemented BMI1, which is expressed by Panc-1, as a key contributor to chemoresistance and invasiveness.^27^ We tested if BCN057 caused an increase in BMI-1 or LGR5 in Panc-1 cells and found no significant increase in the abundance (Figure 5B). In the oncogenic RAS phenotype, PTEN expression is suppressed^24^ developing an antiapoptotic character and we have shown here that inhibition of GSK3β restores PTEN expression and induces apoptosis in this phenotype. As such, our findings are consistent with the suggested reference^26^ as, whether stem like or not, if they are driven by oncogenic RAS, they are susceptible to apoptosis when PTEN is restored. However, on the other hand, in non KRAS mutant stem cells, inhibition of GSK3β has a pro survival effect noted by others^6,35,40^.

Checkpoint inhibitors have transformed the therapeutic space allowing a new arm for intervention for many cancers. Although there is significant progress to be made with combined therapies in this field, the anti-programmed death-1 (PD-1) immune checkpoint inhibitor pembrolizumab (Keytruda®) is the only immunotherapy that is FDA-approved for the treatment of patients with advanced PDAC—provided PDAC is mismatch repair deficient (dMMR) or microsatellite instability high (MSI-H)^41^. However, the objective response rate in solid tumors is relatively low, at best 20 to 30%^42^, and is accompanied by different degrees of side effects. Considering the limited success and challenges of checkpoint blockade in anticancer treatment, new approaches are needed. Here we demonstrate that in an in-vitro co-culture of human T cells and PANC-1 cells, PD-1 expression was significantly reduced in T cells. This is consistent with the known effect of GSK3β inhibition in suppressing PD-1 expression in T-cells. Previous groups^29, 30^, ^31^ have found that GSK3β inhibition resulted in induced expression of T-bet, a repressor of PD-1 gene transcription, which reduces PD-1 expression in CD8+ cytotoxic T cells. We also saw a decrease in PDL-1 expression in PANC-1 cells, however, this was not significant and more likely due to increased apoptotic cell death (as shown in Figures 2 and 4) than regulation of expression. We verified that the antiapoptotic effect of BCN057 on irradiated T-lymphocytes indicates that the decrease in PD-1 expression observed is not due to cell apoptosis (Figure 7B). The EC_50_ value of 5.34 μM is also within the known plasma concentrations achieved by the drug (Supplementary Figure 1). Survival of T-lymphocytes after radiation was also a criteria of the initial screening of the drug^43^.

For both immunotherapy and radiation/chemo therapy intestinal toxicity is a major limiting factor. It is well understood that cells which are rapidly dividing are more susceptible than quiescent cells to radiation and chemotherapy. Rapidly dividing cells also include epithelial tissue stem cells. Because it is known that cytotoxic chemotherapy also targets these rapidly dividing cells, chemotherapy enhances epithelial damage from radiation^44,45^. Thus, intestinal epithelial toxicity is a major limiting factor for effective therapy against pancreatic cancer. We have reported that protection of Lgr5+ve ISCs from radiation treatment induced toxicity is crucial to minimize intestinal epithelial damage^6^. 5FU is a component of FOLFIRINOX and the active metabolized from Capecitabine (Xeloda®) which are commonly used in epithelial and pancreatic cancer therapy. In like fashion, the jejunal sections in Figure 5C clearly showed significant loss of Lgr5+ ISCs in 5FU treated mice. However, as in the radiation case, 5FU treated mice receiving BCN057 demonstrated significant presence of Lgr5+ve ISCs (Figure 5C lower panel) suggesting BCN057 protects ISCs from chemotherapy induced off target effects of on the gastrointestinal epithelium.

In summary our study demonstrates that BCN057 can selectively enhance the cytotoxic effect of chemotherapeutic agents in oncogenic KRAS pancreatic cancer cells. Moreover, it mitigates chemo toxic effects on normal intestinal epithelium suggesting that BCN057 can be considered for use as an adjuvant therapy to improve the therapeutic index. The fact that it can restore apoptosis in the oncogenic RAS phenotype suggests a role in pancreatic cancer treatment. In a previous publication^6^ we demonstrated a model in mice using the syngeneic KRASG13R mutant MC38 tumor cell line^46^ where mice receiving BCN057 had significantly lower tumor burden than control, however, mice receiving BCN057 with radiation treatment had a still lower tumor burden and yet were able to tolerate much higher radiation doses than those receiving radiation alone. This highlights the value of targeting pathways which are wired very differently in the transformed vs non-transformed cells where this dual activity in tumoricidal as well as normal tissue sparing activity may lead to better overall and progression free survival.

## Materials and Methods

### Materials

Human Peripheral Blood Pan-T Cells was purchased from Stem Cell Technologies (Vancouver, BC, Canada) and Panc-1 cells were obtained from the Kansas University Medical Center. 5Fu, Irinotecan, and Oxaliplatin were purchased from Sigma (St. Louis, MO, USA). ImmunoCult-XF T Cell Exp Medium, ImmunoCult HuCD3/CD28/CD2 TCellAct, Hu Recom IL-2 (CHO Expr.) were purchased from Stem Cell Technologies (Vancouver, BC, Canada). The ATPLite Luminescence Assay System and 96-well white, opaque sterile tissue culture treated microplates were purchased from Perkin Elmer (Waltham, Massachusetts, USA). The RealTime Glo Annexin V Apoptosis and Annexin Assay was purchased from Promega (San Luis Obispo, CA, USA). Phospho-GSK-3β (Ser9), PD-L1 (E1L3N). PD-1 (D4W2J), β-Actin, Anti-rabbit IgG, HRP-linked Antibody, Phospho-Akt (T308), Phospho-c-Jun (T93) were purchased from Cell Signaling Technology (Danvers, Massachusetts, USA). 0.4 µm TC Plate Insert with a PC Membrane 6 well plate (thinserts) was purchased from VWR (Radnor Pennsylvania, USA). PD-L1, B7-H1) Monoclonal Antibody MIH1 conjugated with APC and PD-1 Monoclonal Antibody MIH4, conjugated with FITC were purchased from Thermo Fisher Scientific (Waltham, Massachusetts, USA).

Preparation of Compounds: Stock solutions of the following compounds were prepared. 150 mM Oxaliplatin in DMSO, 100 mM Irinotecan in DMSO, 50 mM 5Fu in DMSO, and BCN 057 16 mg/mL in 30% Captisol® (β-cyclodextrin sulfobutyl ether sodium). The vehicle was either DMSO or 30% Captisol® (β-cyclodextrin sulfobutyl ether sodium) without drug.

### Cell lines and cell culture

Panc-1 cells were cultured in DMEM medium supplemented with 10% Fetal Bovine Serum and maintained at 37 °C, 5% CO_2_. Human Peripheral Blood Pan-T Cells were cultured in serum free T cells expansion media with 10 ng/ml of rhIL-2, T-cell activator and expanded as per the manufacturer instructions. Panc-1 cells and T-Cells were grown and expanded separately in T75 flasks. For co-culture experiments with Panc-1 cell, 2.5 million T-cells per well were plated in the well using a 6 well plate. 2.5 million Panc-1 cells were plated in the thinsert which was placed into the well of the 6 well plate in order to have the cells physically separated but share media. The cells were incubated for 24 hrs. at 37 °C, 5% CO_2_. At 24 hrs. the media was replaced with media containing vehicle, or 10 µM BCN057 and were incubated for the following timepoints: 1, 6, 12, 24 hrs. The Panc-1 cells were removed by gently scraping the cells from the thinsert.

### Flow Cytometry

Flow cytometry analysis PANC-1 and T-Cells were harvested independently and washed with 1× PBS. The cells were washed three times with 1× PBS and incubated with antibody for 1 hr. at room temperature. The cells were then washed three times with 1× PBS and resuspended with 1× PBS and analyzed by flow cytometry using a NovoCyte flow cytometer (Agilent Technologies Inc. Santa Clara, CA) and the data was analyzed using the NovoExpress software.

### Western Blot

Cells were harvested and lysed in cell lysis buffer on ice for 15 mins and the lysate was centrifuged at 13,000 rpm at 4 °C for 10 mins. The supernatant was transferred into a new tube for BCA protein quantification assay (Thermo Fisher Scientific) and the samples were normalized. 3x loading buffer (Cell Signaling) was added the protein sample and boiled for 5 minutes. The samples were used for SDS-PAGE analysis and transferred to nitrocellulose membrane. The membrane was blocked with 5% BSA at room temperature for 1 hr. and incubated with primary antibody at 4 °C overnight. The membrane was washed with three times 1x TBST for 5 minutes and incubated with horseradish peroxidase-conjugated secondary antibody for 1 hr. at room temperature. The protein was visualized by SuperSignal West Pico Stable Peroxide Solution (Thermo Fisher Scientific). The blot was visualized by chemiluminescence using a BioRad Versadoc Imaging System. The protein was also visualized using 4-Chloro-1-naphthol solution (Sigma Aldrich) which is a substrate for HRP in a reaction that results in a colored precipitate.

### Viability Assay

To determine the effects of BCN057 in comparison and combination with irinotecan, Oxaliplatin and 5Fu, the ATPLite Luminescence Assay System was used according to manufacturer’s instructions. On day 1, 3,000 Panc-1 cells per well were seeded in 96-well white, opaque sterile tissue culture treated microplates and incubated for 24 hrs. in complete medium at 37 °C, 5% CO_2_. On day 2, the media was exchanged with media containing various combinations of 10 µM BCN 057, 25 µM 5-Fu, 75 µM Oxaliplatin, and 50 µM Irinotecan and incubated at 37 °C, 5% CO_2_. On day 3, 4, and 5, the ATP Lite Assay was performed according to manufacturer’s instructions for the 24, 48, and 72 hrs. timepoints. Unless otherwise stated, values were represented as a percent of the control which is the vehicle for each drug. No difference was found in viability between any control vehicle.

### QPCR

To compare the mRNA levels of BMI1 and LGR5 target genes in Panc-1 cells treated with BCN057 or vehicle. Quantitative PCR (qPCR) was used to measure the expression of the stem cell markers LGR5, and BMI1 which was detected by real-time PCR using the primer pairs listed in Table I. Total RNA was extracted using TRIzol kit (Invitrogen, CA, USA). RNA was reverse transcribed in a final volume of 20 μL using 0.5 μg of oligo dT and 200 U Superscript III RT (Invitrogen) for 30 mins at 50 °C, followed by 2 mins at 94 °C to inactivate the reverse transcriptase. Real-time PCR amplification was carried out in a total volume of 25 μL containing 0.5 μM of each primer, 4 mM MgCl_2_, 12.5 μL of LightCycler™ FastStart DNA Master SYBR green I (Roche Molecular Systems, Alameda, CA) and 10 μL of 1:20 diluted cDNA. PCR reactions were prepared in duplicate and heated to 95°C for 10 mins followed by 40 cycles of denaturation at 95 °C for 15 secs, annealing at 60 °C for 1 min, and extension at 72 °C for 20 secs in ABI Applied Biosystems™ QuantStudio™ Real time PCR machine. Standard curves (cycle threshold values versus template concentration) were prepared for each target gene and for the endogenous reference (GAPDH) in each sample. Quantification of the unknown samples was performed using the nanodrop (Thermofisher scientific).

**Table I:**
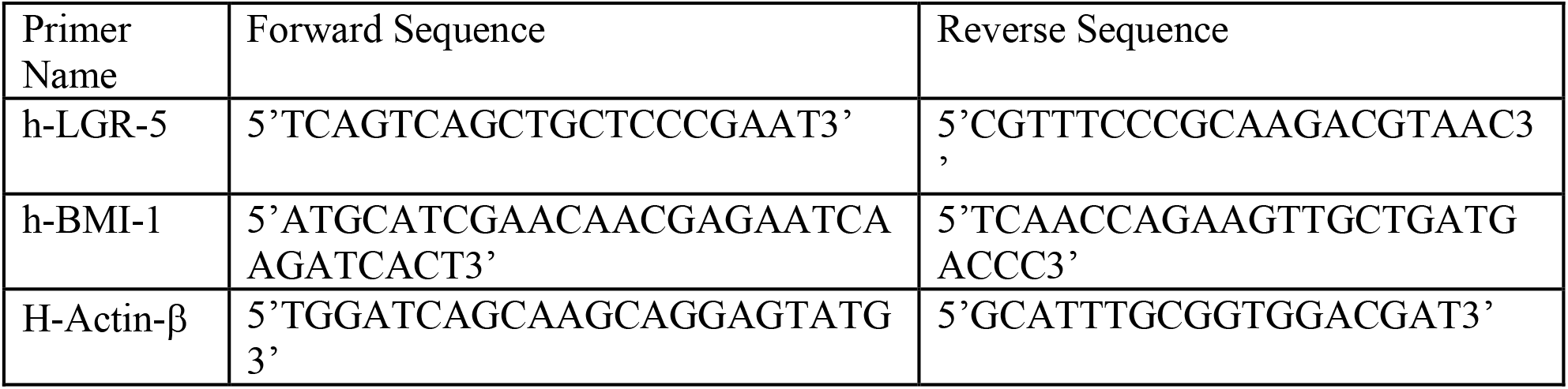
Primers used for qPCR of LGR5 and BMI1 in pancreatic cancer cells Panc-1 treated with 10 µM of BCN057 or vehicle.

### Characterization of BCN057 dose proportionality

Characterization of the proportional plasma exposure vs dose given by subcutaneous (s.c.) administration. Initially, the exposure analysis was performed in three doses: 45, 90 and 150 mg/kg using BCN057 -30% Captisol®, Mass: 401.16 administered via a single s.c. injection in 200 μL. Time points (post dose) were collected by cardiac puncture in euthanized C57Bl/6 mice at 0.08, 0.25, 0.5, 1, 4, 7 and 24 hrs. with 3 mice per time point for a total of 21 animals. Plasma samples (20 μL) were processed by a protein precipitation method. All samples were analyzed with a triple quadrupole mass spectrometer (Agilent^®^ 6460) coupled to an HPLC system (Agilent^®^ 1290) using a reverse-phase analytical column 100 × 2.1 mm Phenomenex Kinetex C18 column (Phenomenex®). The chromatographic conditions consist of mobile phase water and formic acid 0.1% (v/v) (buffer A) and acetonitrile with formic acid 0.1% (v/v) (buffer B) run over a 30-minute gradient elution at a flow rate of 0.15 mL/min with buffer B at 5 % at 5 minutes, 90 % at 20 minutes, 5 % at 21 minutes and 5 % at 30 minutes for equilibration in-between injection cycles. For the analysis of BCN057, 3-[(Furan-2-ylmethyl)-amino]-2-(7-methoxy-2-oxo-1,2-dihydro-quinolin-3-yl)-6-methyl-imidazo[1,2-a]pyridin-1-ium, Exact Mass: 401.16 m/*z;* RT=13.3 min and an internal standard BCN078 which has a fluorobenzyl substitution, Exact Mass: 429.47 m/*z*; RT=13.5 min, the quadrupoles were operated at unit resolution in the positive-ion multiple-reaction monitoring mode (MRM). The MRM experiments were operated with a 20-millisecond dwell time for all transitions (m/z: 401.16→93; 429.47→320). BCN057 was monitored with the transition from m/*z* 401→93 with fragmentor settings at 110 V and a collision energy of 45 and an accelerator voltage of 4 V. BCN 078 was monitored with the transition from m/*z* 429→320 with fragmentor settings at 155 V and a collision energy of 17 and an accelerator voltage of 4 V. Quantitation was performed by use of an external standard method using the analyte BCN057 freebase, yielding a linear regression least-squares fit from 10 pmol to 1000 pmol with R^2^ =0.9983.

### Statistics

The results are shown by calculating the averages (± Standard deviation) in comparison to untreated controls with the indicated (n). The Percent Viability was calculated by dividing the average of each condition by the average of the untreated control (vehicle). The p-values were calculated using the student t-test. Statistical analysis was performed in excel.

## Figures

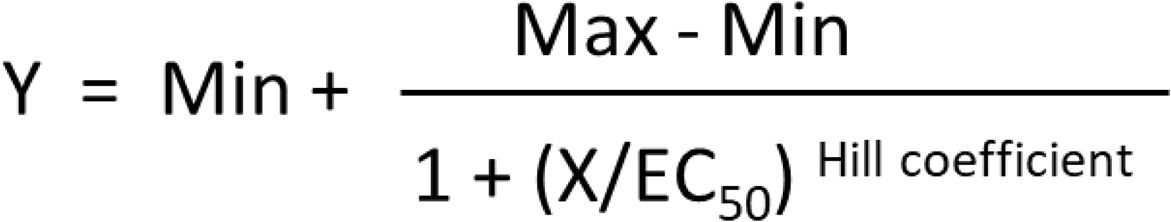

**Equation 1:** Equation representing a four parameter logistic regression model fitting a sigmoid function.

## SUPPLIMENTARY DATA

**Supplementary Figure 1:**
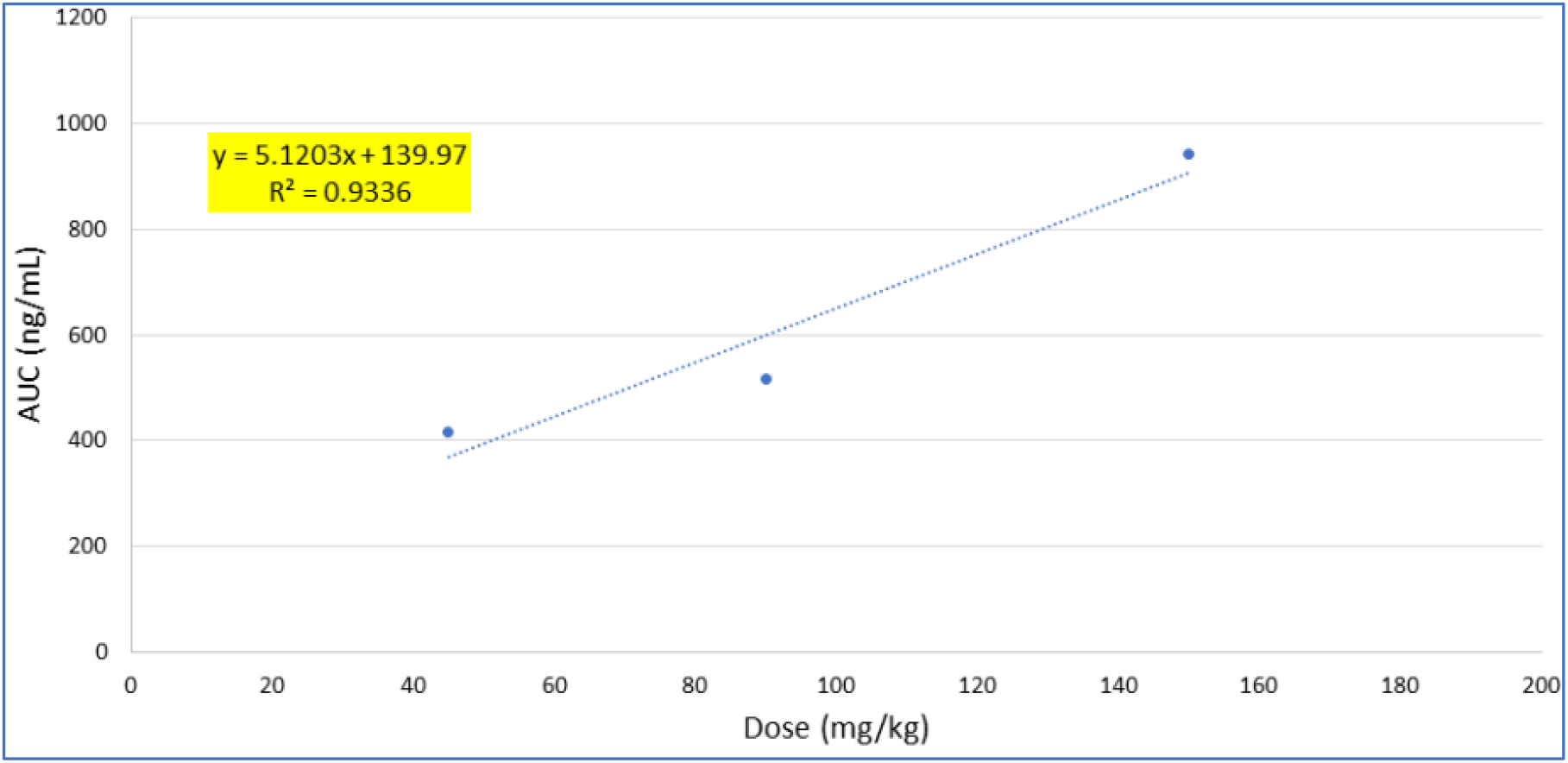
Characterization of BCN057 dose proportionality.

BCN057 - 30% Captisol®, administered via a single subcutaneous (s.c.) injection. Time points (post dose) were collected by cardiac puncture in euthanized C57Bl/6 mice at 0.08, 0.25, 0.5, 1, 4, 7 and 24 hrs with 3 mice per time point. Doses administered were 45, 90 and 150 mg/kg and plotted as a function of AUC (ng/mL) when plotted out to 7 hrs with a least square fit R^2^ = 0.9336. Within this time range, the dose is proportional up to 150 mg/kg when given subcutaneously.

**Supplementary Figure 2:**
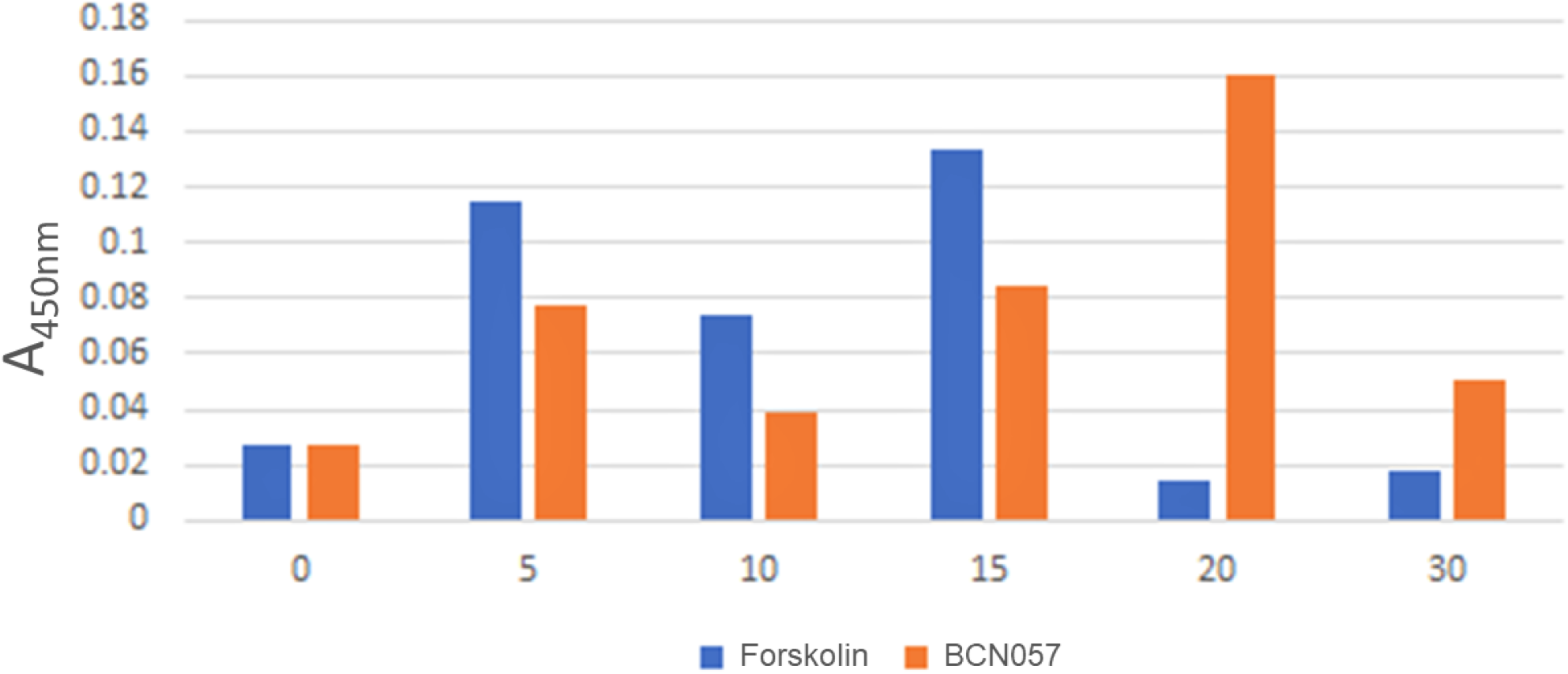
ELISA showing GSK3β phosphorylation at S-9in HEK293 cells at increasing time points after treatment with 10 μM BCN057. Each point is expressed as a ratio of total GSK3β. As a positive control, Forskolin was run as a positive control.

**Supplementary Figure 3:**
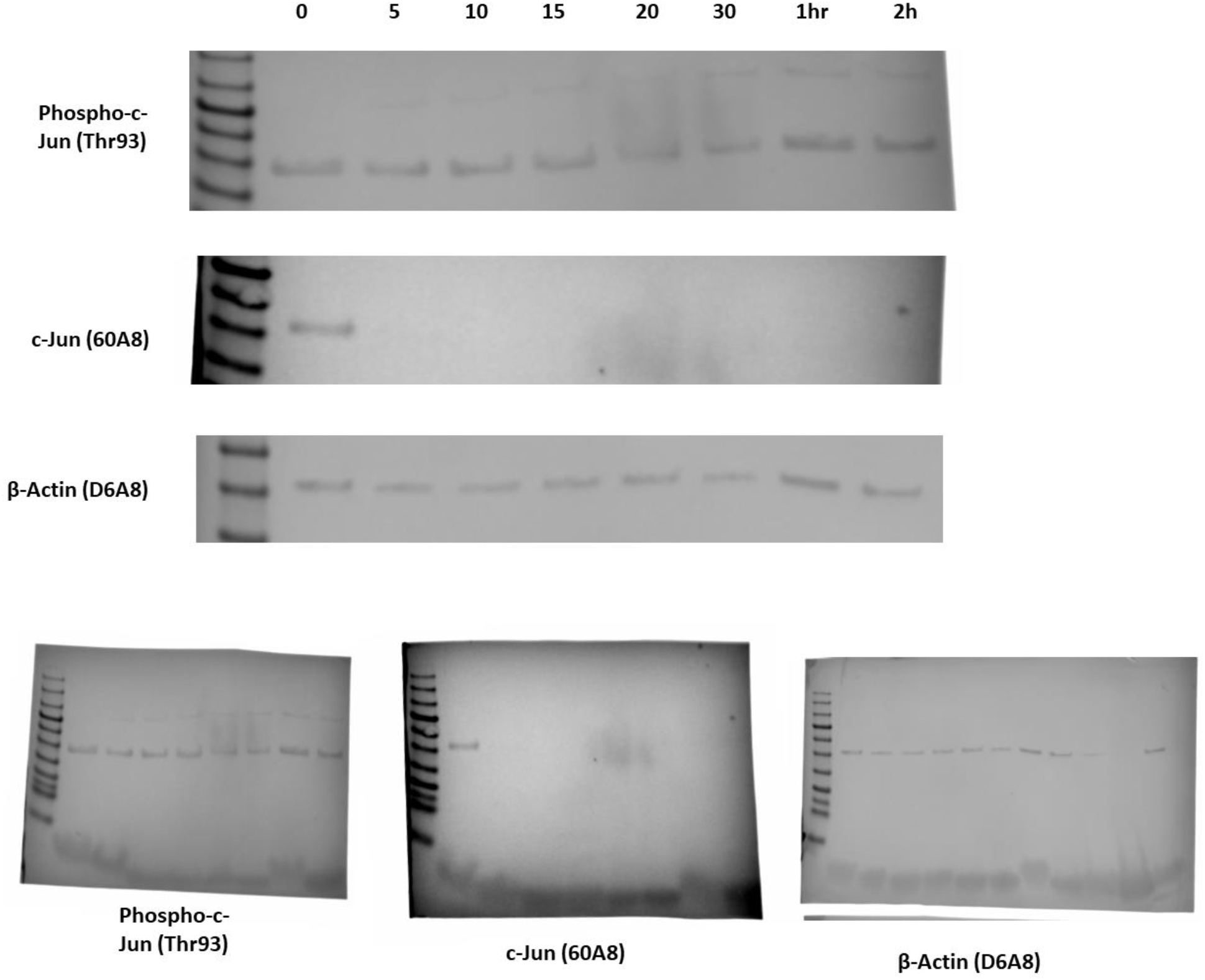
c-Jun Phosphorylation on T93 in HEK293 cells. Non transformed epithelial HEK293 cells were stained for c-Jun phospho T93, a pan specific and β-Actin antibody after treatment with 10 µM BCN057 for up to 2 hrs. No cellular apoptosis or specific T93 phosphorylation state change was observed (First panel). All 3 full western blot gels are shown underneath. However, c-Jun total protein decreased over time (Second panel).

**Supplementary Figure 4:**
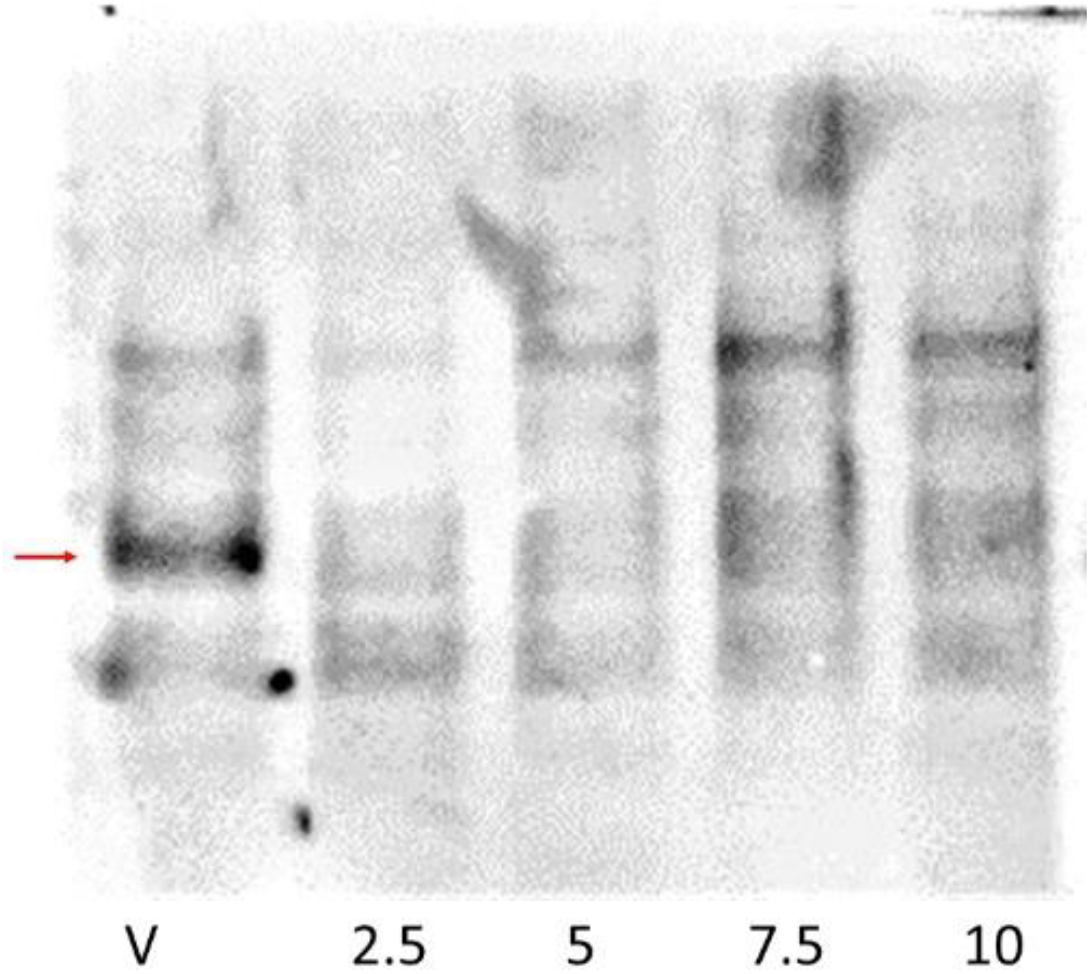
Western blot showing PDL-1 expression (red arrow) on Panc-1 cells co cultured with human T-lymphocytes with no drug (vehicle) or 2.5, 5, 7.5 and 10 hrs. with BCN057.

**Supplementary Figure 5:**
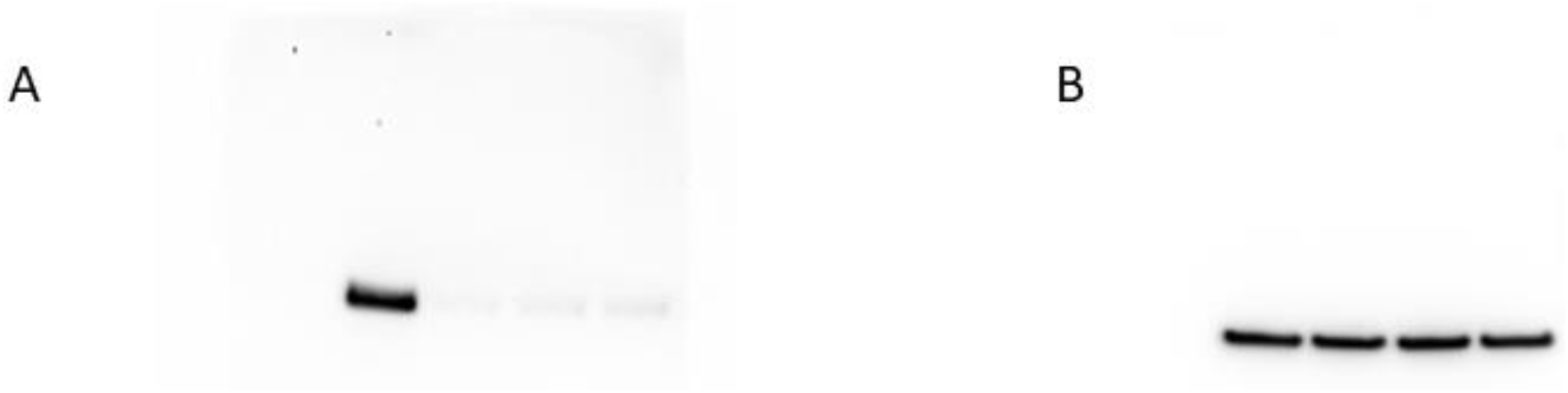
A. Western Blot: PD-1 expression on Human T-lymphocytes co cultured with Panc-1 cells. B. Actin Stain. Actin expression on Human T-lymphocytes co cultured with PANC-1 cells. B. Actin Stain.

**Supplementary Figure 6:**
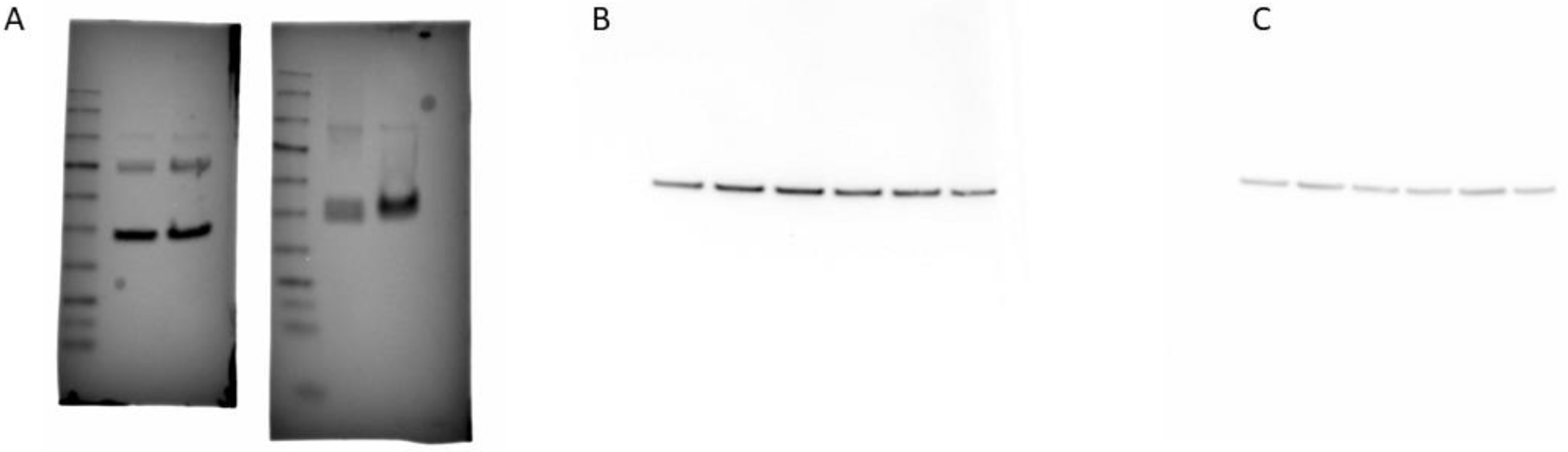
A. c-Jun expression (left panel) and c-Jun T93 phosphorylation in Panc-1 cells treated with 10 µM BCN057. B. PTEN expression at increasing time points as indicated in main manuscript in PANC-1 cells treated with 10 µM BCN057. C. Actin stain from the same blot which has been stripped and re-probed.

**Supplementary Figure 7:**
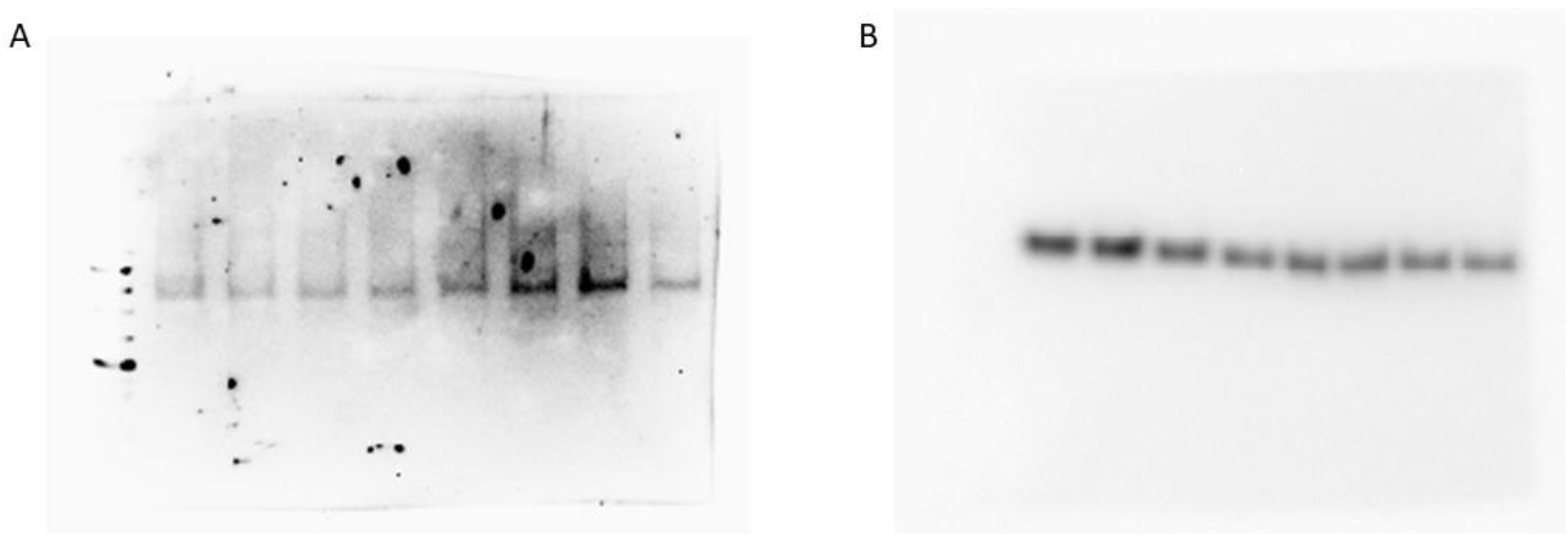
Western Blot Analysis showing, A. GSK3β phosphorylation over time in Panc-1 cells treated with 10 µM BCN057 as described in manuscript. B. Total GSK3β protein expression in the same samples from blot A in PANC-1 cells treated with 10 µM BCN057.

## References

1. Nitsche, C. J., Jamieson, N., Lerch, M. M. & Mayerle, J. v. Drug induced pancreatitis. Best Practice & Research Clinical Gastroenterology 24, (2010).

2. Yadav, D. & Lowenfels, A. B. The Epidemiology of Pancreatitis and Pancreatic Cancer. Gastroenterology 144, (2013).

3. Yang, Q.-J., Zheng, J., Dang, F.-T., Wan, Y.-M. & Yang, J. Acute pancreatitis induced by combination chemotherapy used for the treatment of acute myeloid leukemia. Medicine 99, (2020).

4. American Cancer Society. Key Statistics for Pancreatic Cancer. https://www.cancer.org/cancer/pancreatic-cancer/about/key-statistics.html.

5. Zhan, Y. et al. β-Arrestin1 inhibits chemotherapy-induced intestinal stem cell apoptosis and mucositis. Cell Death & Disease 7, (2016).

6. Bhanja, P., Norris, A., Gupta-Saraf, P., Hoover, A. & Saha, S. BCN057 induces intestinal stem cell repair and mitigates radiation-induced intestinal injury. Stem Cell Research & Therapy 9, (2018).

7. Metcalfe, C., Kljavin, N. M., Ybarra, R. & de Sauvage, F. J. Lgr5+ Stem Cells Are Indispensable for Radiation-Induced Intestinal Regeneration. Cell Stem Cell 14, (2014).

8. Sato, T. et al. Single Lgr5 stem cells build crypt-villus structures in vitro without a mesenchymal niche. Nature 459, (2009).

9. Clevers, H. The Intestinal Crypt, A Prototype Stem Cell Compartment. Cell 154, (2013).

10. Shukla, P. K. et al. Rapid disruption of intestinal epithelial tight junction and barrier dysfunction by ionizing radiation in mouse colon in vivo: protection by N -acetyl-<scp>l</scp> -cysteine. American Journal of Physiology-Gastrointestinal and Liver Physiology 310, (2016).

11. Rahib, L., Wehner, M. R., Matrisian, L. M. & Nead, K. T. Estimated Projection of US Cancer Incidence and Death to 2040. JAMA Network Open 4, (2021).

12. Ikediobi, O. N. et al. Mutation analysis of 24 known cancer genes in the NCI-60 cell line set. Molecular Cancer Therapeutics 5, (2006).

13. Gradiz, R., Silva, H. C., Carvalho, L., Botelho, M. F. & Mota-Pinto, A. MIA PaCa-2 and PANC-1 – pancreas ductal adenocarcinoma cell lines with neuroendocrine differentiation and somatostatin receptors. Scientific Reports 6, (2016).

14. AAT Bioquest, Inc. IC50 Calculator. https://www.aatbio.com/tools/ec50-calculator.

15. Awasthi, N. et al. Comparative benefits of Nab-paclitaxel over gemcitabine or polysorbate-based docetaxel in experimental pancreatic cancer. Carcinogenesis 34, (2013).

16. Giampazolias, E. & Tait, S. W. G. Caspase-independent cell death: An anti-cancer double whammy. Cell Cycle 17, (2018).

17. Ougolkov, A. v., Fernandez-Zapico, M. E., Savoy, D. N., Urrutia, R. A. & Billadeau, D. D. Glycogen Synthase Kinase-3β Participates in Nuclear Factor κB–Mediated Gene Transcription and Cell Survival in Pancreatic Cancer Cells. Cancer Research 65, (2005).

18. Ougolkov, A. v., Bone, N. D., Fernandez-Zapico, M. E., Kay, N. E. & Billadeau, D. D. Inhibition of glycogen synthase kinase-3 activity leads to epigenetic silencing of nuclear factor κB target genes and induction of apoptosis in chronic lymphocytic leukemia B cells. Blood 110, (2007).

19. Reddy, C. E. et al. Multisite phosphorylation of c-Jun at threonine 91/93/95 triggers the onset of c-Jun pro-apoptotic activity in cerebellar granule neurons. Cell Death & Disease 4, (2013).

20. Marchand, B., Tremblay, I., Cagnol, S. & Boucher, M.-J. Inhibition of glycogen synthase kinase-3 activity triggers an apoptotic response in pancreatic cancer cells through JNK-dependent mechanisms. Carcinogenesis 33, (2012).

21. Zhang, J.-S. et al. Mutant K-Ras increases GSK-3β gene expression via an ETS-p300 transcriptional complex in pancreatic cancer. Oncogene 30, (2011).

22. Reddy, C. E. et al. Multisite phosphorylation of c-Jun at threonine 91/93/95 triggers the onset of c-Jun pro-apoptotic activity in cerebellar granule neurons. Cell Death & Disease 4, (2013).

23. Morton, S. A reinvestigation of the multisite phosphorylation of the transcription factor c-Jun. The EMBO Journal 22, (2003).

24. Vasudevan, K. M., Burikhanov, R., Goswami, A. & Rangnekar, V. M. Suppression of PTEN expression is essential for antiapoptosis and cellular transformation by oncogenic Ras. Cancer Research 67, 10343–10350 (2007).

25. Halloran, C. M. et al. 5-Fluorouracil or gemcitabine combined with adenoviral-mediated reintroduction of p16INK4A greatly enhanced cytotoxicity in Panc-1 pancreatic adenocarcinoma cells. The Journal of Gene Medicine 6, (2004).

26. Amsterdam, A., Raanan, C., Schreiber, L., Polin, N. & Givol, D. LGR5 and Nanog identify stem cell signature of pancreas beta cells which initiate pancreatic cancer. Biochemical and Biophysical Research Communications 433, 157–162 (2013).

27. Proctor, E. et al. Bmi1 Enhances Tumorigenicity and Cancer Stem Cell Function in Pancreatic Adenocarcinoma. PLoS ONE 8, (2013).

28. Abrams, S. L. et al. GSK-3β Can Regulate the Sensitivity of MIA-PaCa-2 Pancreatic and MCF-7 Breast Cancer Cells to Chemotherapeutic Drugs, Targeted Therapeutics and Nutraceuticals. Cells 10, (2021).

29. Wu, X. & Witzig, T. E. A new tool in the cancer immunotherapy toolbox? Annals of Lymphoma 1, (2018).

30. Taylor, A., Rothstein, D. & Rudd, C. E. Small-Molecule Inhibition of PD-1 Transcription Is an Effective Alternative to Antibody Blockade in Cancer Therapy. Cancer Research 78, (2018).

31. Li, C.-W. et al. Glycosylation and stabilization of programmed death ligand-1 suppresses T-cell activity. Nature Communications 7, (2016).

32. Neubert, N. J. et al. A Well-Controlled Experimental System to Study Interactions of Cytotoxic T Lymphocytes with Tumor Cells. Frontiers in Immunology 7, (2016).

33. Pottegård, A. et al. Long-term use of lithium and risk of colorectal adenocarcinoma: a nationwide case–control study. British Journal of Cancer 114, (2016).

34. Maeng, Y.-S., Lee, R., Lee, B., Choi, S. & Kim, E. K. Lithium inhibits tumor lymphangiogenesis and metastasis through the inhibition of TGFBIp expression in cancer cells. Scientific Reports 6, (2016).

35. Marchand, B., Tremblay, I., Cagnol, S. & Boucher, M.-J. Inhibition of glycogen synthase kinase-3 activity triggers an apoptotic response in pancreatic cancer cells through JNK-dependent mechanisms. Carcinogenesis 33, (2012).

36. Zhang, J. S. et al. Mutant K-Ras increases GSK-3Β gene expression via an ETS-p300 transcriptional complex in pancreatic cancer. Oncogene 30, 3705–3715 (2011).

37. Kitano, A. et al. Aberrant Glycogen Synthase Kinase 3β Is Involved in Pancreatic Cancer Cell Invasion and Resistance to Therapy. PLoS ONE 8, (2013).

38. Li, J., Mizukami, Y., Zhang, X., Jo, W.-S. & Chung, D. C. Oncogenic K-ras Stimulates Wnt Signaling in Colon Cancer Through Inhibition of GSK-3β. Gastroenterology 128, (2005).

39. Hongisto, V. et al. Lithium Blocks the c-Jun Stress Response and Protects Neurons via Its Action on Glycogen Synthase Kinase 3. Molecular and Cellular Biology 23, (2003).

40. Kazi, A. et al. GSK3 suppression upregulates β-catenin and c-Myc to abrogate KRas-dependent tumors. Nature Communications 9, (2018).

41. Patel, K. et al. Pancreatic Cancer: An Emphasis on Current Perspectives in Immunotherapy. Critical Reviews™ in Oncogenesis 24, (2019).

42. Jiang, Y., Zhao, X., Fu, J. & Wang, H. Progress and Challenges in Precise Treatment of Tumors With PD-1/PD-L1 Blockade. Frontiers in Immunology 11, (2020).

43. Kim, K. et al. High throughput screening of small molecule libraries for modifiers of radiation responses. International Journal of Radiation Biology 87, (2011).

44. Toshkov, I. A. et al. Mitigation of Radiation-Induced Epithelial Damage by the TLR5 Agonist Entolimod in a Mouse Model of Fractionated Head and Neck Irradiation. Radiation Research 187, (2017).

45. Yu J. Intestinal stem cell injury and protection during cancer therapy. Transl Cancer Res. 2, 384–396 (2013).

46. Agalioti, T. et al. Mutant KRAS promotes malignant pleural effusion formation. Nature Communications 8, (2017).

